# Cryopreservation of *Platynereis dumerilii* larvae

**DOI:** 10.1101/2025.07.31.667934

**Authors:** Estefania Paredes, Netsanet Berhane Getachew, Luis Alberto Bezares-Calderón, Sara Campos, Andrij Belokurov, Kristin Tessmar-Raible

## Abstract

The marine annelid *Platynereis dumerilii* is a functional molecular model organism for developmental, evolutionary and chronobiological studies. Research on *Platynereis* is rapidly growing, and with it, the number of genetic variants that laboratories isolate or generate and must subsequently maintain and propagate. Therefore, there is an urgent need to alleviate the burden of live culture maintenance by developing cryopreservation techniques for this species. We report the first cryopreservation protocol for *P. dumerilii* larvae, which combined with a careful post-thawing culturing regime, allowed us to obtain animals that survived to adulthood and successfully reproduced. Our experiments show highest survival rate in 6 – 8 day-old larvae. Equilibration with cryoprotecting agents takes 1h in 5% (v/v) Me2SO + 0.1%(v/w) sucrose, followed by transfer to 0.25ml straws. The protocol cools larvae at 2,5°C/min from 20ºC to -35°C using a programmable freezer, followed by a rapid transfer to liquid N_2_. Larvae are thawed in a water bath at 18°C. The post-thaw larvae feeding regime consisted of 50% *Tetraselmis* + 50 % diatom strains mixture (*Grammatophora marina* and *Nitzschia laevis*). The maximum survival obtained with this protocol so far produced 34% survival after ∼5 months.

## Introduction

The marine annelid *Platynereis dumerilii* has emerged as a significant model organism in chronobiological, neurobiological, developmental and evolutionary biology research. Its unique biological characteristics make it an excellent system for investigating a range of fundamental biological processes, including embryonic development, regeneration, larval biology, lunar and daily rhythms [16,9,18]. Consequently, research using *P. dumerilii* holds considerable potential for advancing our understanding of biological mechanisms. Ensuring continuous access to high-quality biological material is essential for sustaining and expanding research utilizing this model organism.

Animal biobanks play a critical role in scientific research by providing centralized repositories of high-quality biological samples, facilitating efficient and cost-effective studies. These biobanks, whether large-scale international culture collections or small institutional repositories, support a wide range of research endeavors. One of their key advantages is the ability to store and (in some cases) distribute diverse genetic strains and variants, including knockouts and knock-ins, enabling investigations into various biological processes, drug mechanisms, and disease models. However, as biobank collections expand, limitations related to storage space, financial resources, and labor-intensive maintenance present significant challenges.

The cryopreservation and biobanking of model organisms are crucial for the advancement of scientific research. By preserving biological material in a stable, long-term state, researchers can ensure the availability of valuable genetic resources, reducing the dependency on continuous live culture maintenance. This not only minimizes costs and resource consumption but also safeguards against the accidental loss of essential strains due to environmental fluctuations, contamination, or unforeseen disruptions in laboratory operations. Furthermore, cryopreservation allows for the reproducibility of experiments by enabling researchers to access genetically identical samples over time, fostering more consistent and comparable results across studies. The integration of cryopreservation into biobanking efforts is, therefore, a fundamental step toward ensuring the sustainability and reliability of model organism research. Many biobanks house unique strains or variants that have taken years to generate or isolate, yet their long-term preservation is threatened by the high costs associated with maintenance, including supplies, personnel, and facility space. Research on *Platynereis* species is rapidly growing, and with it, the number of genetic variants that laboratories must maintain and propagate. Therefore, there is an urgent need to alleviate the burden of live culture maintenance by developing cryopreservation techniques for this species.

Cryobiology offers a means to store and preserve biological materials for future use. Cryopreservation enables long-term storage, facilitates the transportation of samples across geographic distances, and ensures the availability of rare and valuable biological specimens for research as needed [12]. In the context of polychaete worms, cryopreservation has the potential to establish reference collections for taxonomic and biodiversity studies, as well as to preserve genetic material for molecular research. However, optimizing cryopreservation protocols is crucial to maintaining the viability and integrity of preserved samples.

To date, successful cryopreservation protocols have been established only for the larvae of *Nereis virens* [8], while sperm cryopreservation has been explored for *Arenicola marina* and *Nereis virens* [1] The objective of this study was to develop a cryopreservation protocol for *Platynereis dumerilii* larvae that ensures long-term viability and supports proper post-thaw growth and development, thereby enabling the biobanking of this important model organism.

## Materials and Methods

### Animal Cultures

Animals were kept in a 50/50 mix of natural seawater (NSW, from the North Sea) and artificial sea water (ASW, Tropic Marin Pro Reef Salt), adjusted to a salinity of 35 ppt. pH is maintained between pH 7.9 and 8.2, general culture rooms are maintained at 18-20 °C for optimum growth rate and a 16:8 light/dark cycle. Adults are kept in 6L acrylic boxes containing 1.5 L of ASW and water changes are conducted every 2 weeks. Culture boxes containing juvenile worms less than 2 months of age are not water-changed due to the risk of losing animals and disruption to tube building and growth. *Platynereis dumerilii* worms synchronize their spawning to a moon cycle, mimicked by dim nocturnal lights of different intensities and duration, depending on the exact requirements of the experiments [19,20].

All animal work was conducted according to European guidelines for animal research, and involved animals that have been in continuous lab culture for >60 years. Experiments conducted at ECIMAT marine station were done with larvae supplied from the same parental lines, maintained in the same conditions but the larvae were kept in filtered natural sea water (FSW, 1µm +UVA).

Original Standard feeding regime: new batches are not fed until after they are cultured at 7-13 days old. The original standard diet begins with feeding larvae < 1 month old twice a week with *Tetraselmis marina*. Once they begin forming tubes, they are fed twice a week: the first feeding consists of finely chopped organic spinach (denree Blattspinat or similar), and the second feeding consists of PlanktoMarin (a diatom-based liquid food, GROTECH GmBH) or, alternatively, a mixture of powdered Tetramine (Tetra) and *Spirulina spec*.

Improved feeding regime: in further experiments that ultimately resulted in successful survival (up to maturing worms) the starter food for the thawed larvae was modified such that only 50% of the mixture consists of *Tetraselmis marina*, the other 50% are 1:1 mixture of diatoms *Grammatophora marina* (Lyngbye) Kützing, 1844 and *Nitzschia laevis* (Nitzschia) Hassall, 1845. Initially, f/2 (Guillard and Ryther 1962, Guillard 1975) was used for growing diatoms. However, we subsequently started using a set of commercial components from CORALAXY. 6-day cultures with a final concentration of ∼1.6–1.8 ⨯ 10 □ cells/ml are used for feeding.

In addition to this live algal diet fed twice a week, larvae were also fed once per week with the commercial food Planktomarin (GROTECH GmBH). After the worms began forming tubes, they were fed with approximately one small chopped organic spinach leaf per worm.

### Cryoprotectant toxicity assays

Cryoprotectant agents (CPAs) used along the experiments were dimethyl sulfoxide (Me_2_SO), ethylene glycol (EG), propylene glycol (PG), Glycerol (Gly), Sucrose (SUC) from Sigma Aldrich. To test the sensitivity to different CPAs, larvae were exposed for 3 minutes to 1.4M of each permeating CPA (Me2SO, EG, PG and Gly), then carefully filtered (40 µm mesh) and transferred to clean sea water for observation. In a second phase, combinations of cryoprotecting agents were tested by reducing Me2SO content and supplementing it with glycerol and/or sucrose and proceeded as above.

### Cryopreservation

Cooling was done using a controlled rate freezer Cryologic (Australia, LLD) or Consarctic BIOFREEZE BV65/88. The cryopreservation protocol with the highest efficiency according to Olive and Wang [8] for their target species consisted of loading 0.25 ml straws (IMV technologies), with a 1:1 mix of larvae and different concentrations of Me2SO. Larvae were incubated for ca. 3,10,30 and 60 min in the cryoprotecting solution. Straws were sealed with colored sealing powder (IMV technologies) and cooled down from 20°C to -35°C at 2.5min ^-1^, Straws were transferred immediately into liquid nitrogen. Thawing of each straw was done in a water bath at 18°C by immersion for 20-30 seconds. Thawed larvae were rinsed in filtered NSW. The morphology of the larvae was documented before and after each trial with a stereomicroscope, and in some cases with a compound microscope. Mortality rates were obtained from counts of damaged and seemingly healthy larvae.

### Histology

Histological analysis was done by the Estación de Ciencias Mariñas de Toralla (ECIMAT) service using haematoxylin and eosin (H&E), which is a common staining protocol for polychaete histology, staining nuclei blue and cytoplasm pink [7].

### Statistical analysis

Data were analysed with the non-parametric test Kruskal-Wallis using the SPSS (version 15.0), p < 0.05 was regarded as significant.

## Results

The initial protocol tested was based on the protocol of Olive and Wang [8] on 8 days post-fertilization (dpf) *Nereis virens* larvae. It consisted of 1.4M Me2SO for 3 minutes, cooling at 0.3ºC/min to -25ºC than 2.5ºC/min to -35ºC and liquid nitrogen transfer. We used this protocol as our reference and applied it on *Platynereis dumerilii* larvae of the same age using 1.4M PG, EG or Me2SO. We first ensured that none of these cryoprotectant agents (CPAs) had deleterious effects on animal survival. Table 1 shows that a 3 min exposure to any of these CPAs has no effect on larval mortality even a week post exposure. We therefore included the three CPAs in our initial cryopreservation trials. A high initial survival was obtained (80, 63.9 and 60.8% larvae survived) immediately after thawing when incubated for 3 min in Me2SO, EG or PG respectively. By day 5, survivors were found only in treatments cryopreserved in Me2SO, but the larvae looked lethargic and moved only spasmodically. A day later, survival dropped to 39% and to 0% by day 8 (Table 1). Surviving larvae presented notorious damage in the posterior (pygidium) area (Figure 1 A-C) that worsened with time.

**Table 1.** Eight-days old larvae were incubated in a CPA solution for 3 minutes in straws at 20ºC. Then they were cryopreserved and thawed using a slow rate freezer, following Olive and Wang (1997) protocol. Maximum mOsmols of solutions (SW 1000+ CPA1400) =2400 mOsmol. N=100 larvae per straw and the experiment was replicated 3 times with WildType (WT) polychaetes, results show average ±SD.

| Cryoprotectant | Exposure time (min) | Cryopreserved (Y/N) | Preliminary exploratory survival (n=1) immediately post-exposure/thaw (%) | Survival Day 5 (n=3) exposure/post-thaw (%) | Survival day 6 (n=3) post exposure/-thaw(%) | Survival day 8 (n=3 post exposure/thaw (%) |
| --- | --- | --- | --- | --- | --- | --- |
| Me2SO 1.4M | 3 | N | 100 | n.a. | 100 | n.a. |
| EG 1.4M | 3 | N | 100 | n.a. | 100 | n.a. |
| PG 1.4M | 3 | N | 100 | n.a. | 100 | n.a. |
| Me2SO 1.4M | 3 | Y | 80.0 | 89.9±11.7 | 39.1±17.7 | 0 |
| EG 1.4M | 3 | Y | 63.9 | 0 | 0 | 0 |
| PG 1.4M | 3 | Y | 60.8 | 0 | 0 | 0 |

**Figure 1.**
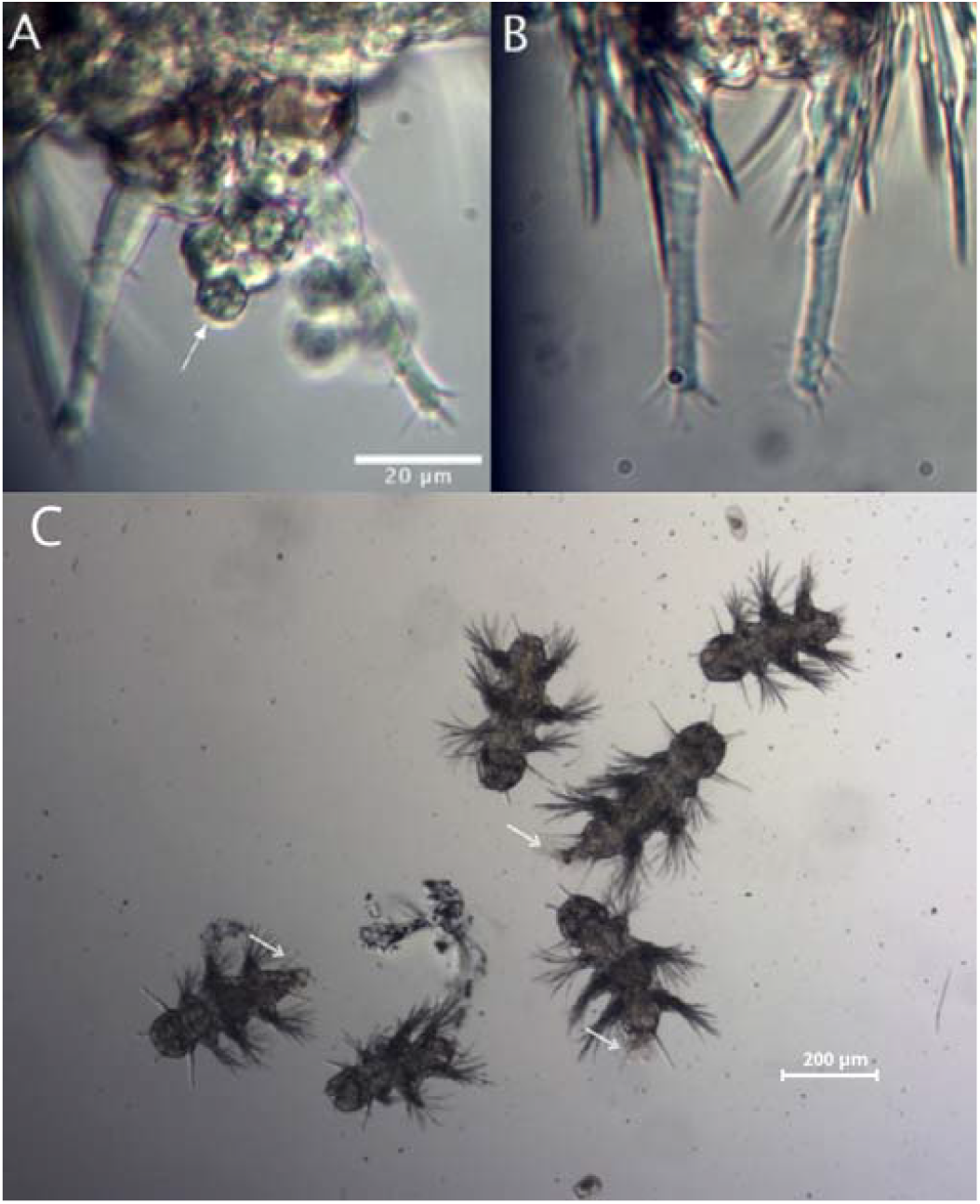
Documentation of initial survival and tissue damage: (A) *P. dumerilii* 6-8 dpf Larvae survived thawing after being cryopreserved according to protocol by Olive and Wang (1997). Close up view of the damaged area, tissue extrusion pointed by arrow. (B) Undamaged control larvae. (C) Larvae showed damaged in the posterior area (white arrows).

We surmised that the tissue damage observed could be in part due to incomplete penetration of the CPAs. To explore this possibility, increasing equilibration times were tested before cooling. Survival rates did not decrease with equilibration times up to 30 minutes (Figure 2). In agreement with our hypothesis, the immediate post-thaw survival was higher the longer the CPA exposure, from 18.7% at a 60 min-equilibration to almost no survival when equilibration times were lower than 10 minutes (Figure 2).

**Figure 2.**
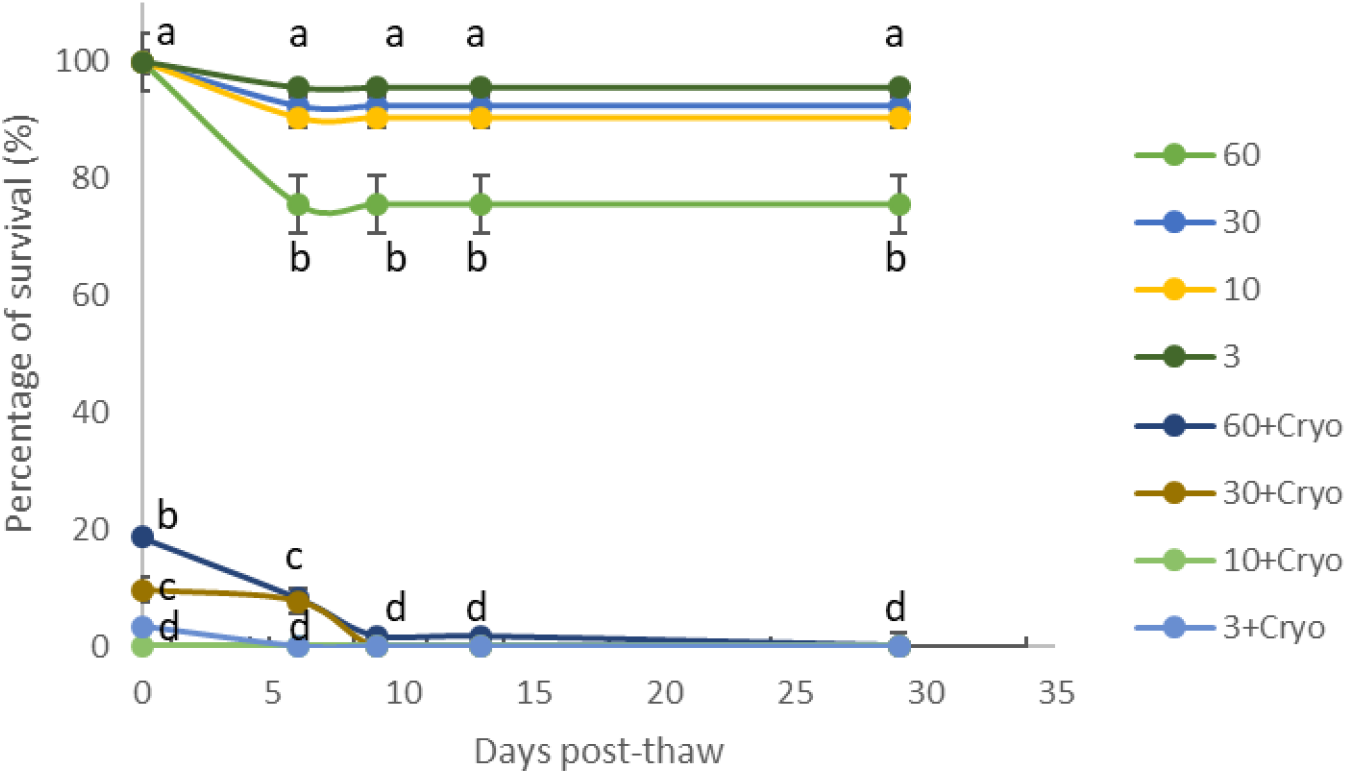
Percentage of survival ±SD after 1.4M Me2SO exposure and exposure plus cryopreservation of 8 dpf larvae at four equilibration times, 3, 10, 30 and 60 min with n=100 larva. The experiment was replicated 3 times with wild-type (wt) animals. Letters highlight statistically significant differences between treatments at the same time slots among all treatments, p>0.05.

CPAs have different modes of action, some penetrating the tissue while others protecting the surface. Therefore, we next tested combinations of CPAs that could provide a more integral protection effect. We specifically tested lower concentrations of Me2SO supplemented with other CPAs. We monitored survival for 29 days with 30 min equilibration time in each of the CPA mixtures. This approach resulted in increased survival rates (Table 2). Specifically, with 0.7M Me2SO+0.03M Suc 28% larvae survived by day 29. Me2SO supplemented with Gly 0.68M and 0.1% sucrose resulted in 47.2% survival (29 days post-thaw, table 2). However, all larvae were dead by day 29 post-thaw. Finally, all larvae died when incubated in 0.68M glycerol and 0.8M Me2SO. In summary, reducing the Me2SO concentration to the half, from 1.4M to 0.7M, extended the post-thaw survival from day 6 (Table 1) to day 13 (Table 2).

**Table 2.**
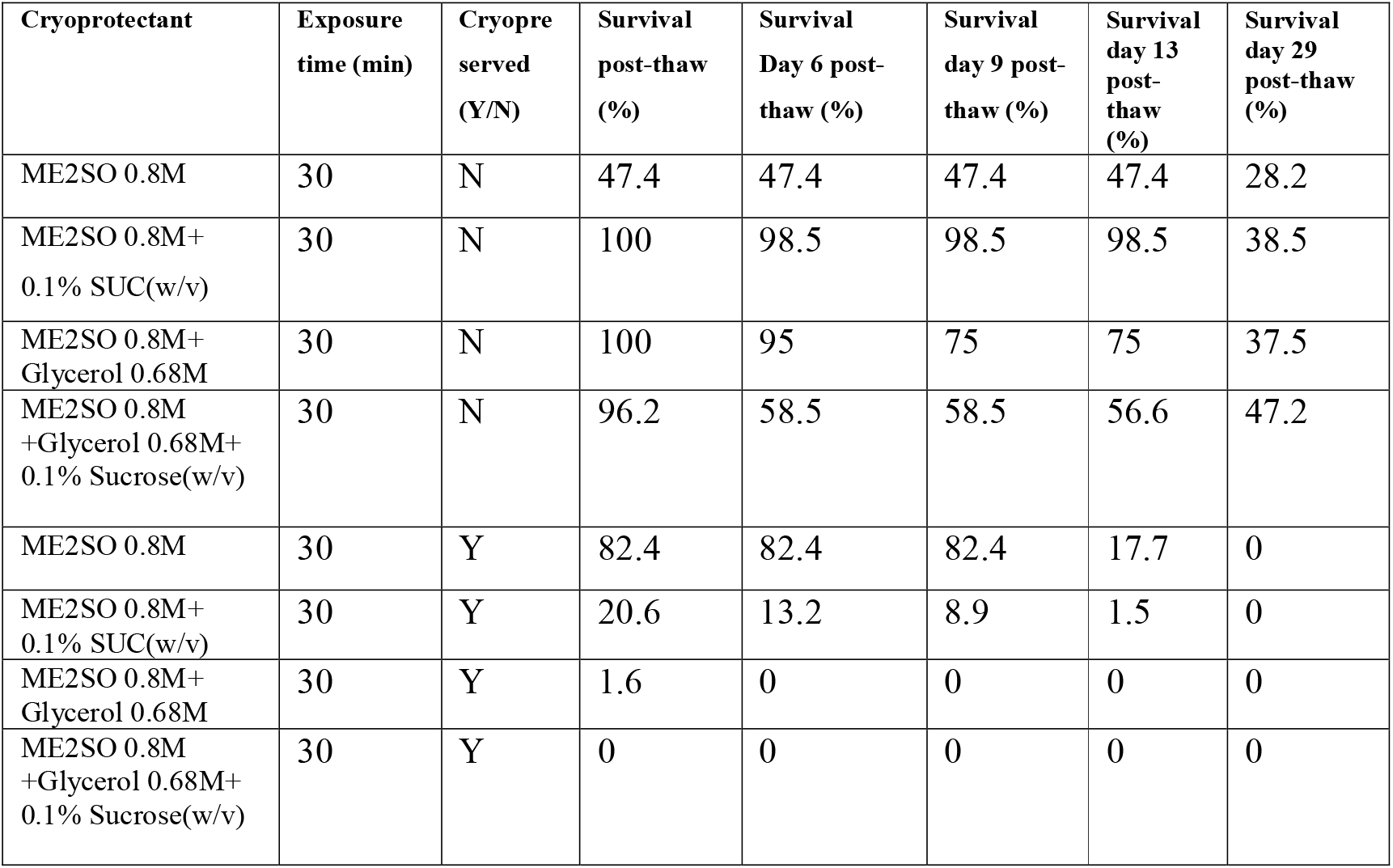
Larvae (8 days old) were incubated in different CPAs for 30 minutes in straws at 20ºC. Then they were cryopreserved using a slow rate freezer, descending the temperature 2.5ºC/min until reaching -35ºC. Next they were transferred into liquid nitrogen. For thawing, a 18ºC bath was used. Maximum mOsmols of solutions (SW 1000+ CPA) = from 1800 to 2480 mOsmol.

Based on these results, a new protocol was developed with the lower Me2SO concentration, 5%Me_2_SO(v/v)+0.1% Sucrose (w/v) (0.7M Me2SO + 0.03M Sucrose), equilibration time increase to 30 min to 1 hour. Cooling directly from 20^a^C to -35ºC at 2.5ºC/min and then transferred to liquid nitrogen. Thawing at a lower temperature of 18ºC water bath. Results shown in figure 3 show that with these adjustments survival could be extended to 57 days post-thaw.

**Figure 3.**
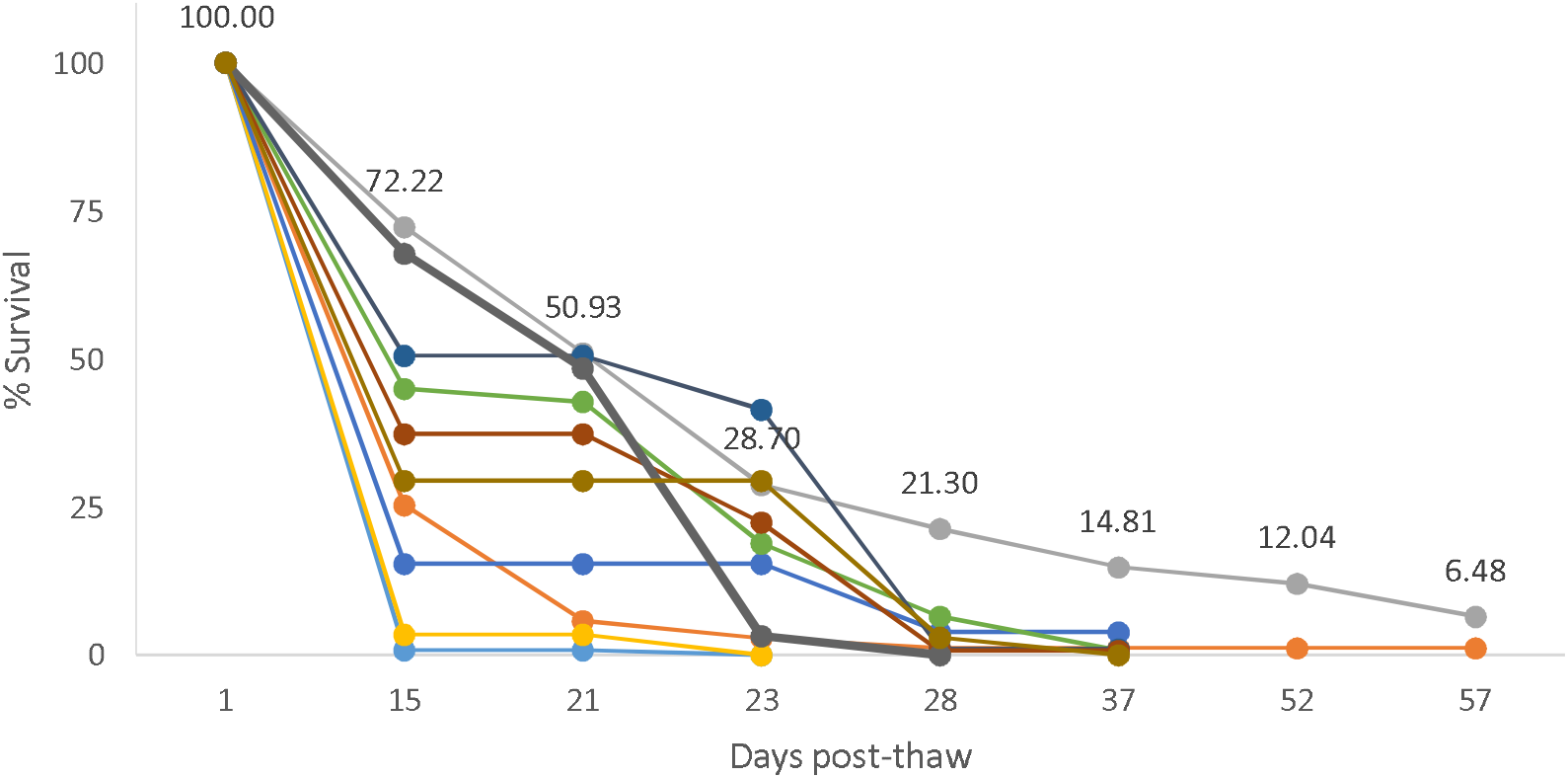
Percentage of survival after increased equilibration time and CPA mixture. Each line represents larvae from a different spawning event followed along time post-thaw, n (individuals) variable between 26 and 176 between batches. Two sets showed survival up to 57 days post thaw, numbers reflect the survival of the best batch obtained. Cryoprotecting solution consists on Me_2_SO 5%+0.1% sucrose, equilibration time 1 hour, cooling directly from 20ºC to -35ºC at 2.5ºC/min and thawing at 18ºC water bath.

This new protocol using 5%Me2SO+0.1% Suc and a lower thawing temperature was tested with 10 different batches of polychaetes and the damage observed in the cryopreservation experiments using the published protocol (Table 1, Figure 1) was no longer present. There were also no external differences in morphology between controls and cryopreserved larvae. To verify this result at the histological level, we analyzed 50 individual histological cuts. Examining the structure of the cryopreserved larvae showed no major internal disruption of the tissues, including the gut and pygidium areas (Figure 4). Worms subjected to this protocol, survived up to 50-60 days (Figure 3) although low in numbers (1-6.5%).

**Figure 4.**
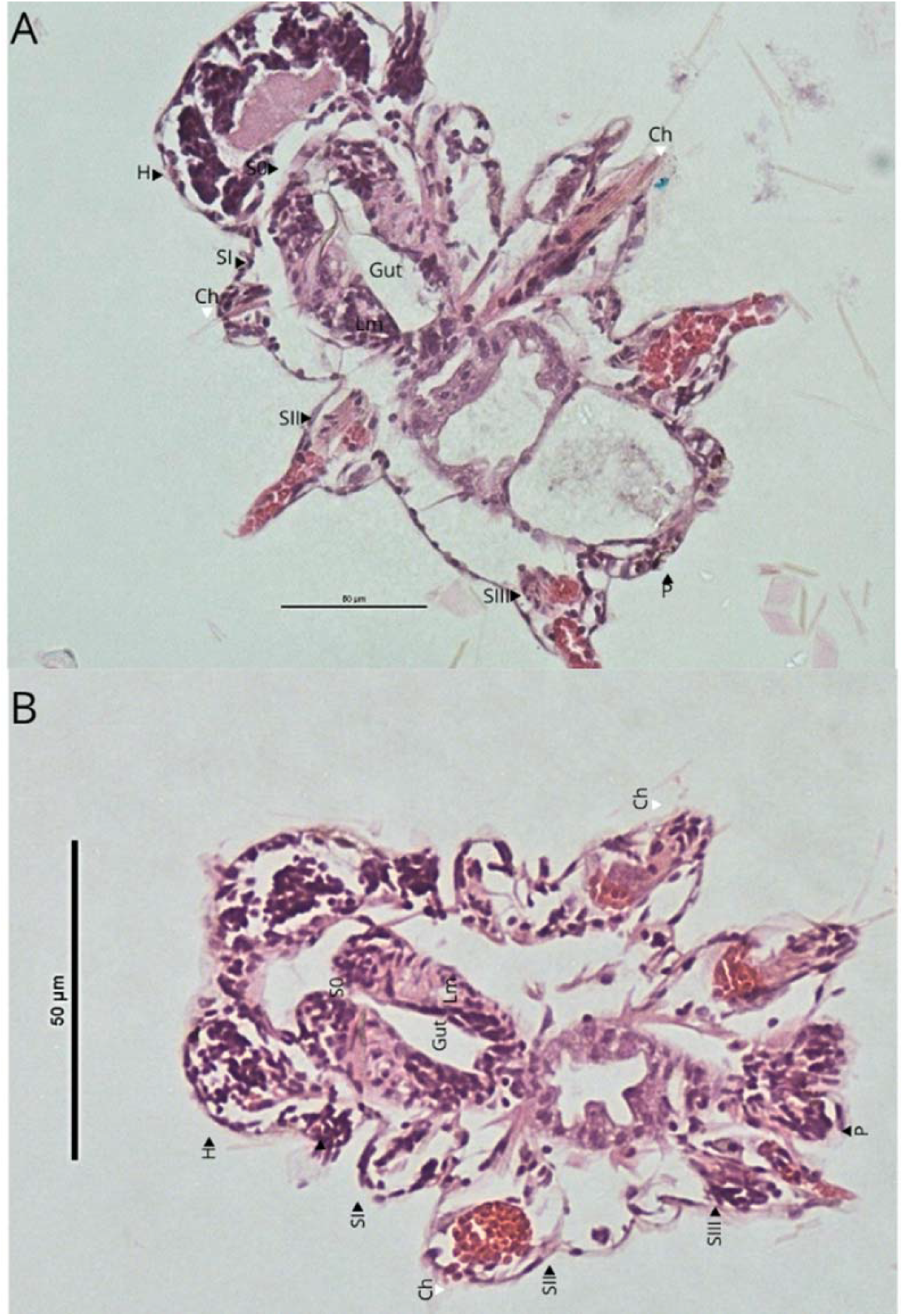
Histological transversal sections of two *P. dumerilii* larvae. (A) Control larvae with a normal morphology at the pygidium (P) area and (B) cryopreserved larvae showing normal tissue in the pygidium (P) area. H (head), Segment zero (S0), First segment (SI), Second segment (SII) and Third segment (SIII). In some segments chaetae can be seen (Ch), Gut and Lm (lateral muscle), the pygidium area (P). Images analyzed n=50

We attributed the gradual reduction in survivor worms to an overall weak health after thawing. Weaker worms could be more susceptible to a suboptimal diet and water quality. Therefore, we modified the feeding regime of the survivor larvae to an improved and more closely monitored diet (see animal culture in Methods: standard vs. improved feeding regime). In brief, the feeding regime builds on recent discoveries on the settlement of *Platynereis* larvae and consisted of an green algal (*Tetraselmis marina*) and diatom (*Grammatophora* and *Nitzschia*) mixture once per week, and once per week of the commercial food Planktomarin [3]. After the worms began forming tubes, they were fed a small amount of finely chopped organic spinach (see materials and methods for further details). We minimized the risk of microbial contamination due to the presence of sucrose by making sure to thoroughly wash the larvae prior to long-term culture, thereby diluting the CPA mix, as well as constantly monitoring water quality of the culture boxes.

This improved post-thaw feeding and caring regime consistently expanded the lifespan of the cryopreserved larvae and led to replicable and significantly improved survival (Figure 5). The average lifetime of the larvae handled by the optimized protocol (see flowchart Figure 7) is >8 months, with polychaetes grown from cryopreserved larvae reaching adult stage with many new segments (Figure 6A) and able to form a tube (Figure 6B). Some of the survivors have also successfully matured (both males and females) (Figure 8). Although the surviving worms grow well and with an average rate that fluctuates from 4.5 to 34%, the cryopreservation treatment may delay the onset of maturation

**Figure 5.**
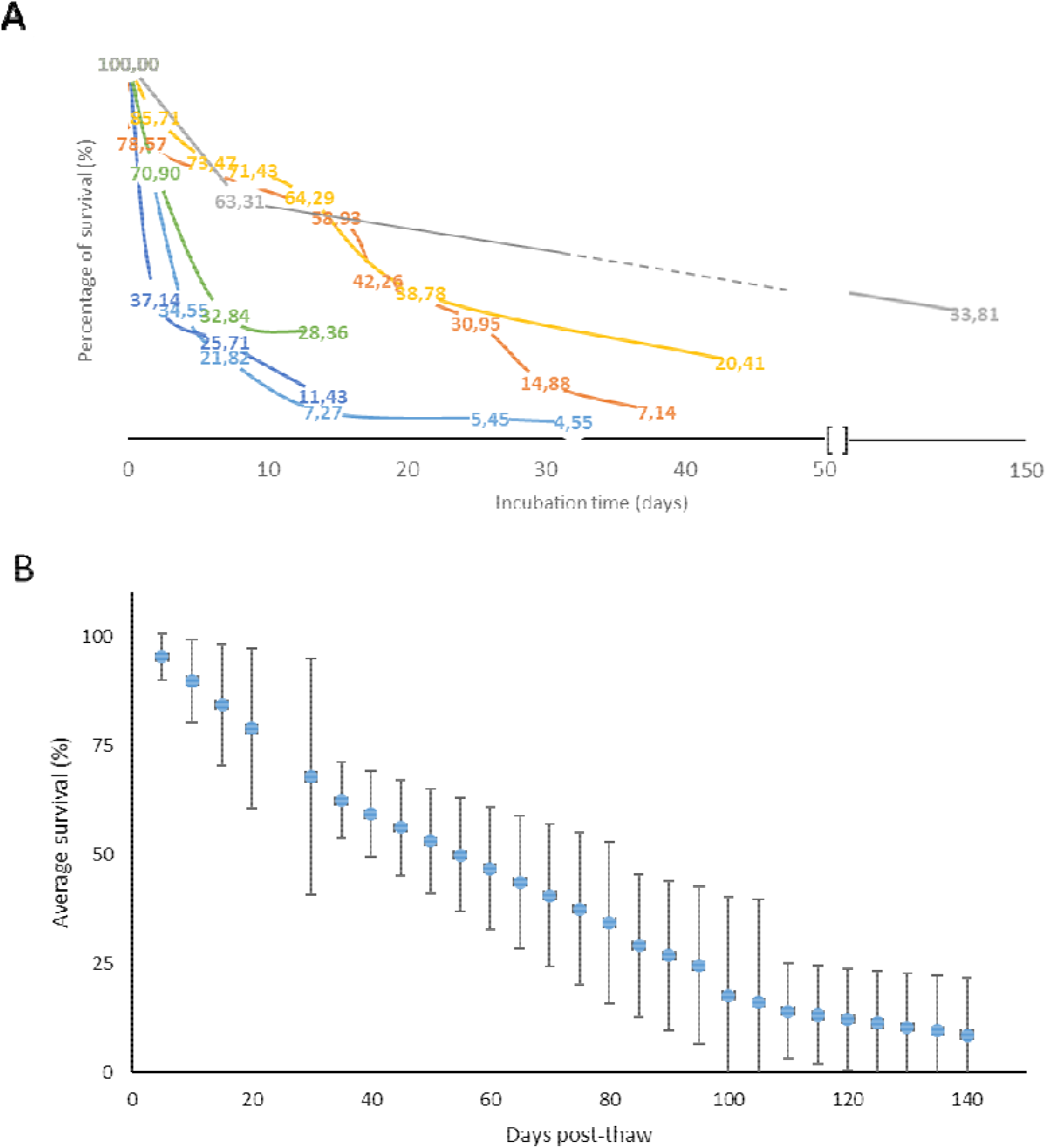
Overview on survival rate and age. The same data are plotted as two different representations.to obtain a better overview on the variance between different batches (A), but also a general average (B). (A) Percentage of survival for the finalized cryopreservation protocol. Each line is a replicate set from a different original spawning event, n variable between 35 and 168, as a function of time. (B) Averaged survival percentage across time after standardizing the incubation time and after each batch was referred to their control. Average ±SD

**Figure 6.**
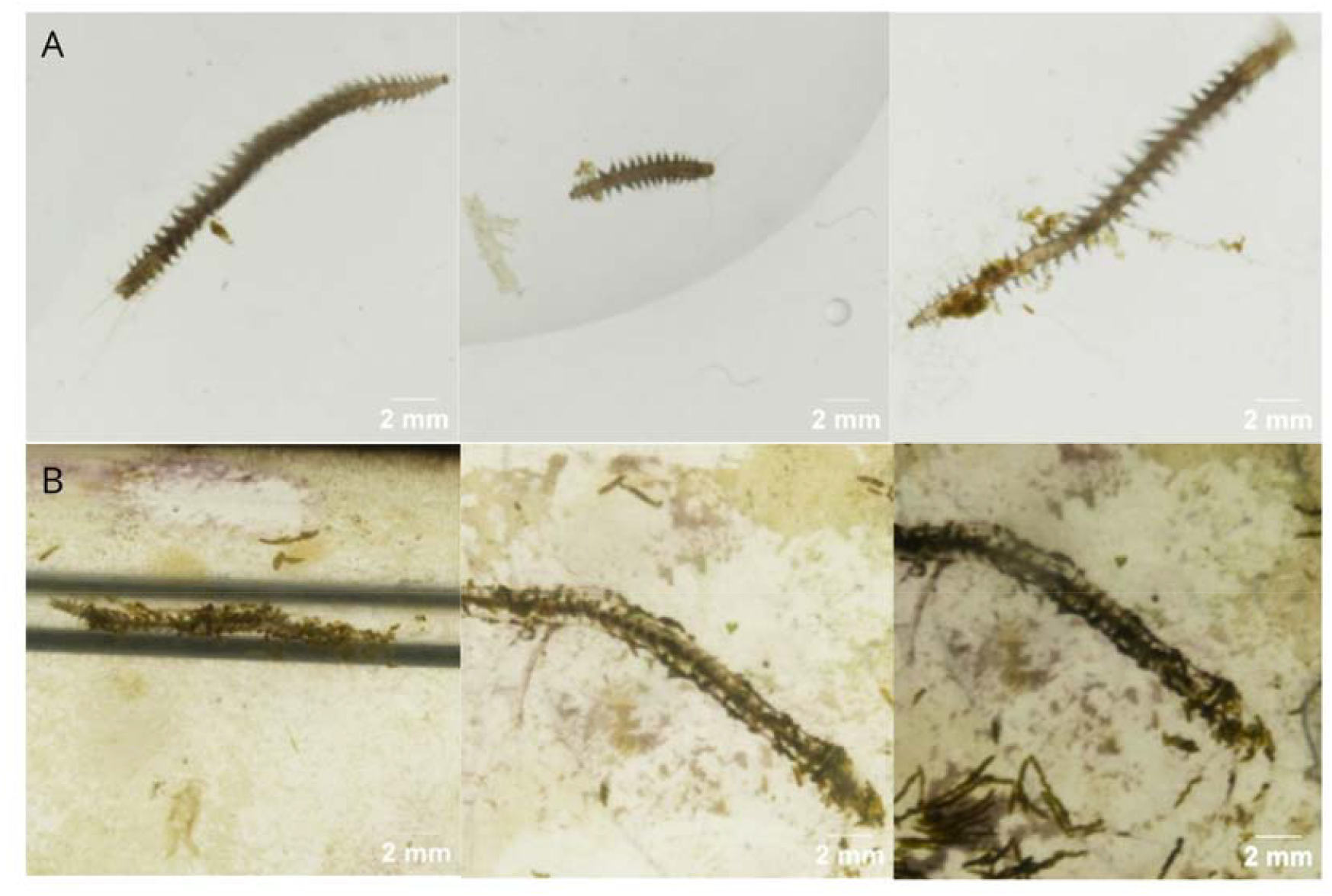
*Platynereis dumerilii* grown from cryopreserved larvae. Many new segments have been added (A) and tubes are normally formed (B).

**Figure 7.**
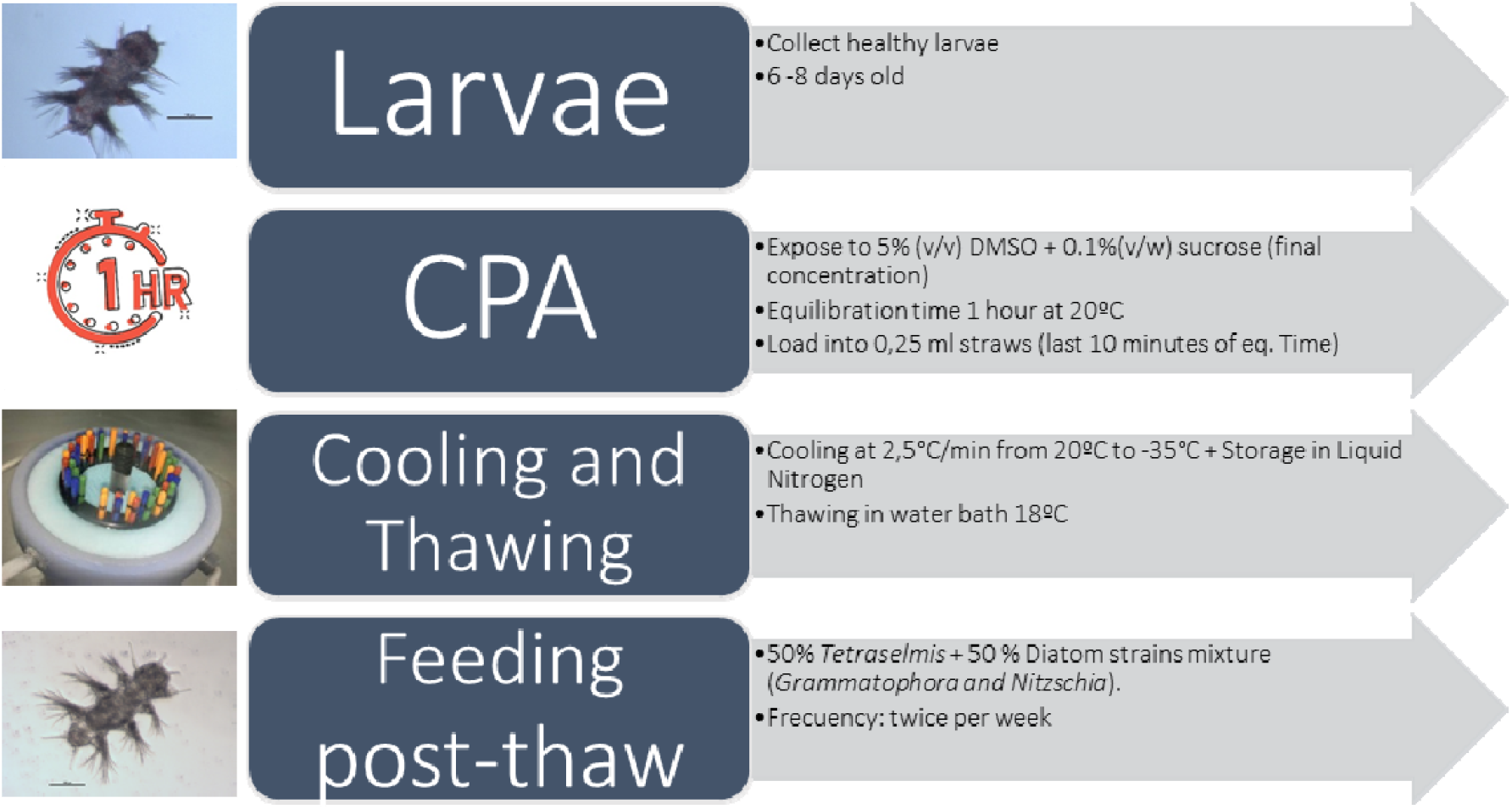
Flowchart with the summary of the successful cryopreservation protocol steps for *P. dumerilii*.

**Figure 8.**
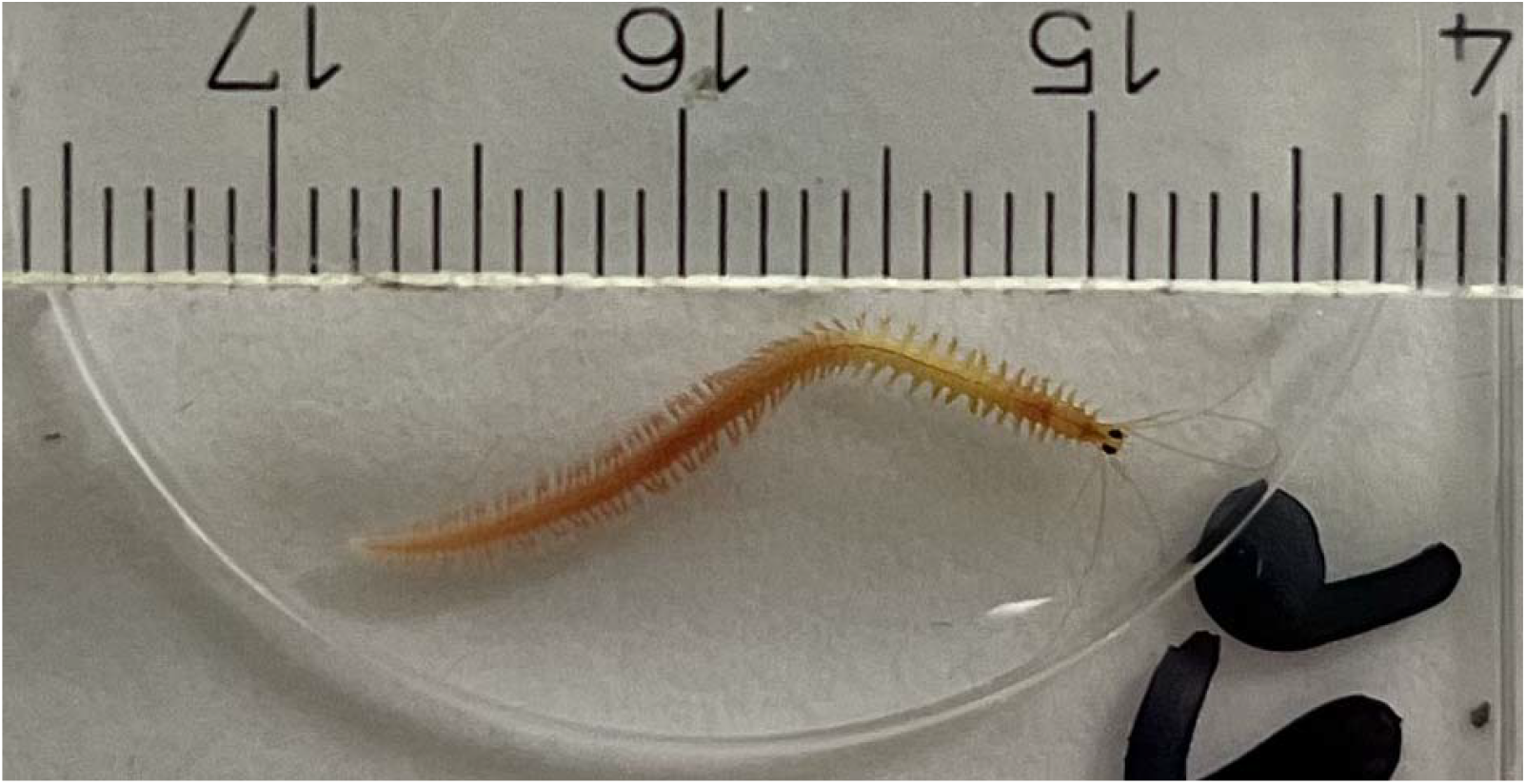
Worms successfully mature after cryopreservation. Epitoke male (4.5 months old) grown from cryopreserved larvae, which gave later rise to surviving offspring. Scale in cm.

## Discussion

The objective of this study was to develop a cryopreservation protocol for *Platynereis dumerilii* larvae that ensures long-term viability and supports normal post-thaw growth and development. We developed a protocol that allows the cryopreservation of 6 to 8 dpf larvae with a maximum survival rate of 33.8% 150 days post-thawing. Surviving individuals are normal in all measured aspects: they form segments and can form silk tubes that they use as shelter. Some of the survivors have attain sexual maturity, albeit at a slower rate than worms in our reference culture. This is the first cryopreservation protocol established for this polychaete species, and only the second for polychaetes, following that developed for *Nereis virens* [8].

These results underscore the importance of tailoring cryopreservation protocols to specific species, particularly through detailed assessments of cryoprotectant (CPA) toxicity [10,14,2]. When a protocol exists for a closely related species, it may serve as a valuable starting point; however, as demonstrated here, species-specific modifications—such as adjustments to cooling or warming rates are often necessary [13]. One effective strategy to mitigate CPA toxicity involves the use of combined cryoprotectants that provide enhanced cryoprotection with reduced toxicity. Typically, a mixture of permeant and non-permeant CPAs is employed [12]. In the case of *P. dumerilii*, the inclusion of low concentrations of sucrose in Me2SO led to an increased survival rate immediately after thawing and alleviated the damaging effects of the protocol during post-thaw culture. The protective role of sucrose likely stems from its ability to stabilize cellular membranes during osmotic transitions and reduce intracellular ice formation [15].

Moreover, reduction of the thawing bath temperature from 35°C to 18°C likely contributed to alleviate the previously observed physical damage localized in the pygidium region in the protocol developed by Olive and Wang [17]. Although rapid warming is generally associated with improved survival outcomes [6,11], in some cases, thawing at temperatures closer to the organism’s physiological range yields better results, as reported for other marine invertebrates [10] by reducing the uneven warming and risk of overheating. Finally, once the protocol was producing a reliable survival percentage post-thaw the final obstacle was to maintain the larvae incubated post-thaw with the maximum level of fitness. By applying the new feeding regimen proposed by Hird, et al [3], the cryopreserved larvae were able to develop further and reach a sexually mature stage, establishing the effectiveness of the developed protocol.

This improvement of cryopreservation outcomes with a change of feeding regime highlights the importance of the larvae fitness. This results are in agreement with data recently found in other marine invertebrate larvae whose post-thaw survival improved when being fed even before cryopreservation but also post-thaw [4,5]. *P*. dumerilii is increasingly used as a molecular model in evolutionary developmental biology, neuro- and chronobiology. The availability of a cryopreservation method enhances laboratory flexibility by decoupling breeding from experimental demand and maintaining genetic consistency across experiments. Post-thaw developmental competence, including the ability to grow new segments and achieve reproductive maturity, suggests that cryoinjury is minimal in essential somatic and germline tissues.

## Acknowledgements

E.P Holds a Grant Ramon y Cajal grant by MICIU/AEI/10.13039/501100011033 and by “European Union NextGenerationEU/PRTR”. L.A.B.C received Transnational Access funds from Assemble+, grant from the European Union’s Horizon 2020 research and innovation programme (No. 730984). KTR has been supported by the Helmholtz Society, distinguished professorship by the Alfred Wegener Institute Helmholtz Centre for Polar and Marine Research, H2020 European Research Council, ERC Grant Agreement #819952, Austrian Science Funds (FWF), SFB F78 and CoE Neuroscience, Human Frontier Science Program (HFSP), #RGP021/2024. None of the funding bodies was involved in the design of the study, the collection, analysis, and interpretation of data or in writing the manuscript. Authors would like to thank technical staff from both the ECIMAT marine station in Spain and the University of Vienna aquatic infrastructure for expert animal maintenance and care.

## Declaration of interests

The authors declare no competing interests.

## Notes

### Competing Interest Statement

The authors have declared no competing interest.

### Summary of Updates

An author name was misspelled and needed corrected. Nothing else has changed.

## References

1. Bury, N.R., & Olive, P.J.W. Ultrastructural observations on membrane changes associated with cryopreserved spermatozoa of two polychaete species and subsequent mobility induced by quinacrine. Invertebrate Reproduction & Development 23, 139–150 (1993). 10.1080/07924259.1993.9672305

2. Heres, P., Rodriguez-Riveiro, R., Troncoso, J., & Paredes, E. Toxicity tests of cryoprotecting agents for Mytilus galloprovincialis (Lamark, 1819) early developmental stages. Cryobiology 86, 40–46 (2019). 10.1016/j.cryobiol.2019.01.001

3. Hird, C., Jékely, G., & Williams, E.A. Microalgal biofilm induces larval settlement in the model marine worm Platynereis dumerilii. Royal Society Open Science 11, 240274 (2024). 10.1098/rsos.240274

4. Lago, A., & Paredes, E. Modulation of stress factors for cryopreservation of Paracentrotus lividus (Lamarck 1816) larvae. Cryobiology 110, 8–17 (2023). 10.1016/j.cryobiol.2023.01.006

5. Lago, A., Troncoso, J. & Paredes, E. Cryopreservation of juvenile Mytilus galloprovincialis to safeguard mollusk biodiversity and support aquaculture. Sci Rep 15, 25587 (2025). 10.1038/s41598-025-11439-3

6. Mazur, P. Principles of Cryobiology. In: Fuller, B.J., Lane, N., & Benson, E.E. (eds) Life in the Frozen State, 3–65. CRC Press, Boca Raton (2004). 10.1201/9780203647073.ch1

7. Murray, H.M., Gallardi, D., Gidge, Y.S., Sheppard, G.L. Histology and Mucous Histochemistry of the Integument and Body Wall of a Marine Polychaete Worm, Ophryotrocha n. sp. (Annelida: Dorvilleidae) Associated with Steelhead Trout Cage Sites on the South Coast of Newfoundland. Journal of Marine Dciences, (2012). 10.1155/2012/202515

8. Olive, P.J.W., & Wang, W.B. Cryopreservation of Nereis virens (Polychaeta, Annelida) larvae: The mechanism of cryopreservation of a differentiated metazoan. Cryobiology 34, 284–294 (1997).

9. Özpolat, B.D., Randel, N., Williams, E.A., et al. The nereid on the rise: Platynereis as a model system. EvoDevo 12, 10 (2021). 10.1186/s13227-021-00180-3

10. Paredes, E., & Bellas, J. Cryopreservation of the sea urchin Paracentrotus lividus embryos: Preliminary studies. Cryobiology 59, 179–181 (2009). 10.1016/j.cryobiol.2009.09.010

11. Paredes, E., & Mazur, P. The survival of mouse oocytes shows little or no correlation with the vitrification or freezing of the external medium, but the ability of the medium to vitrify is affected by its solute concentration and by the cooling rate. Cryobiology 67, 386–390 (2013). 10.1016/j.cryobiol.2013.09.003

12. Paredes, E., Heres, P., Anjos, C., & Cabrita, E. Cryopreservation of marine invertebrates: From sperm to complex larval stages. In: Wolkers, W.F., & Oldenhof, H. (eds) Cryopreservation and Freeze-Drying Protocols. Methods in Molecular Biology 2180, 289–307. Humana, New York, NY (2021). 10.1007/978-1-0716-0783-1_18

13. Paredes, E., Campos, S., Lago, A., Bueno, T., Constensoux, J., & Costas, D. Handling, reproducing and cryopreserving five European sea urchins (Echinodermata, Klein, 1778) for biodiversity conservation purposes. Animals 12, 3161 (2022). 10.3390/ani12223161

14. Smith, J.F., Adams, S.L., Gale, S.L. et al. Cryopreservation of Greenshell™ mussel (Perna canaliculus) sperm. I. Establishment of freezing protocol. Aquaculture 334–337, 199–204 (2012). 10.1016/j.aquaculture.2011.12.027

15. Strauss, G., & Hauser, H. Stabilization of lipid bilayer vesicles by sucrose during freezing. Proceedings of the National Academy of Sciences of the United States of America 83, 2422–2426 (1986). 10.1073/pnas.83.8.2422

16. Veedin Rajan, V.B., Häfker, N.S., Arboleda, E., et al. Seasonal variation in UVA light drives hormonal and behavioural changes in a marine annelid via a ciliary opsin. Nature Ecology & Evolution 5, 204–218 (2021). 10.1038/s41559-020-01356-1

17. Wang, W.B., & Olive, P.J.W. Effects of cryopreservation on the ultrastructural anatomy of 3-segment Nereis virens larvae (Polychaeta). Invertebrate Biology 119, (v3) (2005). 10.1111/j.1744-7410.2000.tb00017.x

18. Wulf, P.O., Häfker, N.S., Hofmann, K., & Tessmar-Raible, K. Guiding light: Mechanisms and adjustments of environmental light interpretation with insights from Platynereis dumerilii and other selected examples. Zoological Science 42, 52–59 (2025). 10.2108/zs240099

19. Zantke, J., Ishikawa-Fujiwara, T., Arboleda, E., Lohs, C., Schipany, K., Hallay, N., Straw, A. D., Todo, T., & Tessmar-Raible, K. Circadian and circalunar clock interactions in a marine annelid. Cell reports, 5(1), 99–113 (2013). 10.1016/j.celrep.2013.08.031

20. Zurl, M., Poehn, B., Rieger, D., Krishnan, S., Rokvic, D., Veedin Rajan, V. B., Gerrard, E., Schlichting, M., Orel, L., Ćorić, A., Lucas, R. J., Wolf, E., Helfrich-Förster, C., Raible, F., & Tessmar-Raible, K. Two light sensors decode moonlight versus sunlight to adjust a plastic circadian/circalunidian clock to moon phase. Proceedings of the National Academy of Sciences of the United States of America, 119(22), e2115725119 (2022). 10.1073/pnas.2115725119

